# Probing the effect of uniaxial compression on cell migration

**DOI:** 10.1101/082461

**Authors:** Nishit Srivastava, Robert R Kay, Alexandre J Kabla

## Abstract

The chemical, physical and mechanical properties of the extra-cellular environment have a strong effect on cell migration. Aspects such as pore-size or stiffness of the matrix influence the selection of the mechanism used by cells to propel themselves, including pseudopod or blebbing. How a cell perceives its environment, and how such a cue triggers a change in behaviour are largely unknown, but mechanics is likely to be involved. Because mechanical conditions are often controlled by modifying the composition of the environment, separating chemical and physical contributions is difficult and requires multiple controls. Here we propose a simple method to impose a mechanical compression on individual cells without altering the composition of the gel. Live imaging during compression provides accurate information about the cell’s morphology and migratory phenotype. Using *Dictyostelium* as a model, we observe that a compression of the order of 500 Pa flattens the cells under gel by up to 50%. This uniaxial compression directly triggers a transition in the mode of migration, from primarily pseudopodial to bleb driven, in less than 30 sec. This novel device is therefore capable of influencing cell migration in real time and offers a convenient approach to systematically study mechanotransduction in confined environments.

## Introduction

Cell migration is an important part of both healthy and pathological biological processes. During embryo development, wound healing or immune response, cells have to navigate through complex environments to shape tissues or perform their physiological function (Martin, 1997; Miller and Davidson, 2013; Bonnans et al., 2014). Cell migration is also a defining feature of cancer metastasis Friedl and Gilmour, 2009). However, any understanding of *in vivo* cell migration relies on our ability to study how cells perceive and respond to the chemical and mechanical cues from the environment (Eyckmans et al., 2011). It is already well established that many biochemical cues directly influence the behaviour of a cell (Alberts et al., 2002), but mechanical aspects are only starting to be well understood. The specific properties of the mechanical environment, in particular its stiffness and confining effect, directly influences single cell migration and collective invasion of cells (Bergert et al., 2012; Vedula et al., 2012; Tozluoglu et al., 2013; Wolf et al., 2013; Paluch and Raz, 2013). Cells can sense a variety of external mechanical signals that interact with proteins such as integrin, paxillin and other signaling molecules in cytoplasm like Rho and Rho-associated kinases (Chen et al., 2004; Lämmermann and Sixt, 2009; Gardel et al., 2010; DuFort et al., 2011; Abu Shah and Keren, 2013; Charras and Sahai, 2014). There is however a lack of quantitative understanding of how different mechanotransductive pathways operate in a cell in order to sense mechanical information and control as a result key biophysical quantities such as membrane tension, membrane-cortex attachment or actin polymerisation that regulate cell migration (Houk et al., 2012; Tyson et al., 2014). A simple experimental model environment with tunable mechanical characteristics would enable a systematic study of cells’ migratory response to mechanical stimuli.

Current experimental techniques to study cell mechanics include local methods such as optical or magnetic tweezers (Neuman and Nagy, 2008; De Vlaminck and Dekker, 2012), AFM cantilevers (Bufi et al., 2015), micropipette aspiration (Dai et al., 1999). These techniques offer a convenient way to impose a force on individual cells, but are not well suited to the study of migratory cells which would rapidly move away from the probe. Microfluidic systems and microfabrication techniques are popular approaches to study the migration of cells in different geometries and mechanical environments (Liu et al., 2015; Raab et al., 2016; Denais et al., 2016), on different topographical patterns (Kim et al., 2009) and micropillars (Ghassemi et al., 2012; Lam et al., 2012). These techniques have been instrumental in our understanding of the forces and molecular mechanisms required for migration in confined spaces, probing in particular the role of nucleus during cell migration (Raab et al., 2016). In the case of confinement, the environment inside such devices is however much stiffer than most single cells and soft tissues, whose stiffness is typically in the range of 0.1-10 kPa (Discher et al., 2005). While these are ideal techniques to probe the mechanisms of migration inside preexisting geometrical gaps, it does not fully mimic the broad class of situations where a cell has to create its path through surrounding tissues (Charras and Sahai, 2014).

Under-agarose assays offer an alternative route to study cell migration and mechanotransduction closer to physiological conditions, in particular in conjunction with cell chemotaxis (Nelson et al., 1975; Laevsky and Knecht, 2001). This technique has been utilized for studying the migration of neutrophils and *Dictyostelium discoideum* cells, and allowed an exploration of the key molecular pathways involved in chemical sensing (Kay et al., 2008; Nichols et al., 2015). In such assays, the mechanical properties of the hydrogel can be tuned to study its effect on the cell migration. Zatulovskiy et al. have shown that *Dictyostelium discoideum* cells switch from pseudopodial mode of migration to bleb mode when the stiffness of the hydrogel is increased. In such experiments, modulation of the stiffness is achieved by changing the gel concentration, and hence the pore size and chemical composition of the environment (Normand et al., 2000). It is therefore possible that the stiffness is not a primary parameter controlling the mode of cell migration during such under-agarose assays. Rather than tuning the stiffness of the gel, a mechanical load could however be used to modulate the mechanical environment around the cells. King et al. for instance applied known weights on a slab of agarose gel to probe the role of pressure on autophagy in *Dictyostelium*. In this paper, we extend this approach and build an experimental system which is designed to dynamically impose a mechanical load on a cell migrating under an agarose layer. Since load and gel concentration are controlled independently, the contributions of the matrix stiffness (or composition) and load can be separated. Specifically, we show that a compressive load on *Dictyostelium* can be used to control the mode of cell migration under agarose and open the door to a systematic study of the transduction pathways involved.

## Device design

The primary goal of the device, referred to as *cell squasher* in this article, is to apply a steady and uniform compressive stress on a slab of hydrogel while simultaneously performing high resolution live imaging of cells squashed between the gel and a glass coverslip. The overall design of the device is shown in Figure 1. A rectangular plunger (perspex, typically 4 mm wide, 10 mm length, 3 mm thickness) is used to compress the upper surface of the gel. The vertical position of the plunger is controlled using a motorised translational stage (Newport, TRA-25CC, range 25 mm) so that the load can be dynamically controlled. The pressure imposed by the plunger on the gel is measured by a tension-compression
load cell. The horizontal position of the plunger relative to the hydrogel can be adjusted with two manually controlled linear stages.

**Figure 1:**
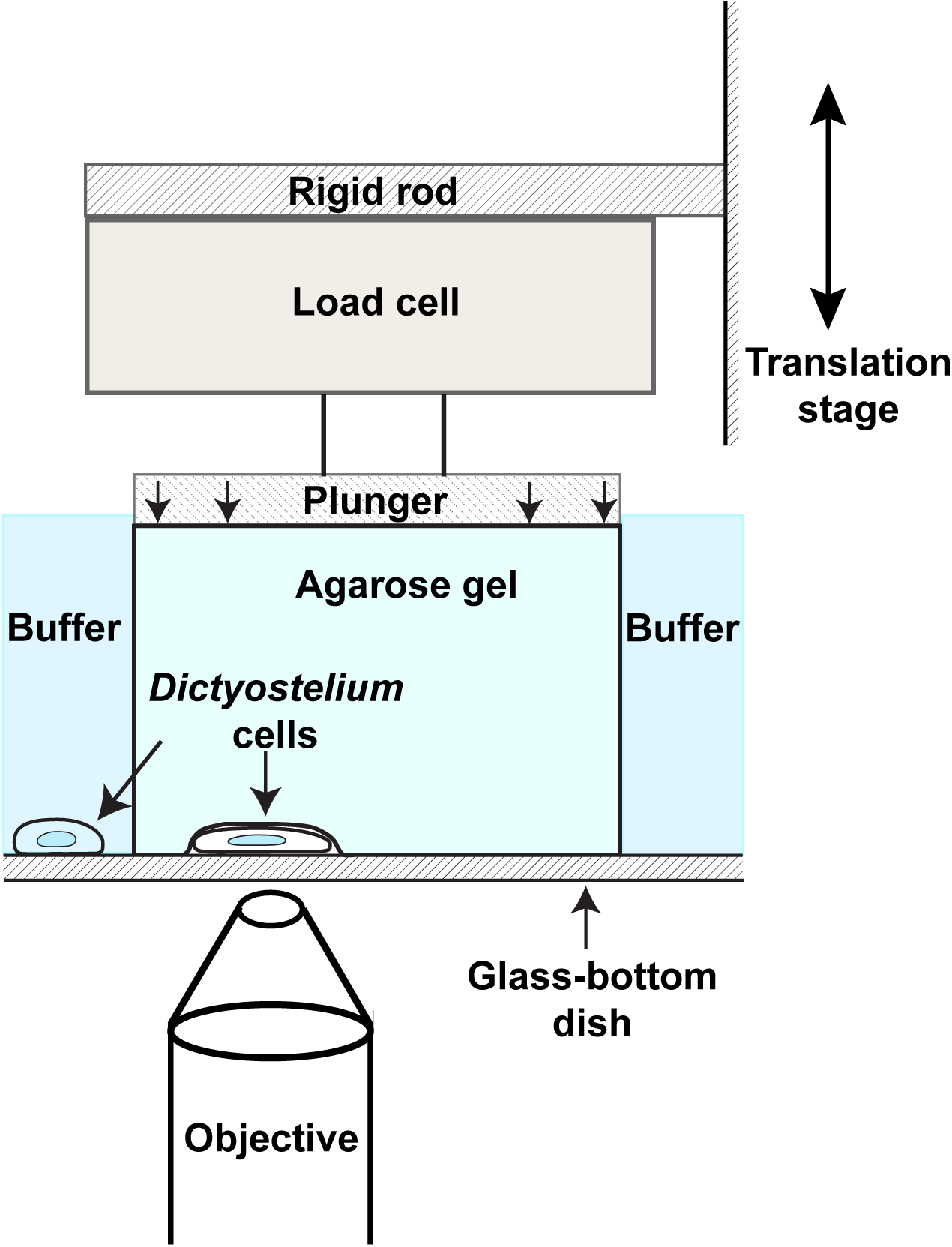
Working principle of the cell squasher. A mechanical load is applied uniformly on a hydrogel while cells are migrating underneath the gel in a classical under-agarose assay. The plunger’s vertical position is controlled by an automated translation stage. The pressure applied is monitored with a load cell feeding back to the stage control system to ensure an accurate and dynamic control of the loading conditions.

The plunger, load cell and positioning system with its motorised actuator needs to reside on the stage of the microscope (Zeiss LSM780, 160 mm length, 110 mm width) so that both move together as a combined unit while selecting regions to image. The stage can bear loads upto 60 N. As a result, the cell squasher is designed to be as compact as possible (121.9 mm length, 133.3 mm width, 95.2 mm height), making the device fairly portable and usable on a broad range of inverted fluorescence microscopes. The load cell-plunger system furthermore needs to be accommodated between the condensor and lens of the microscope (20 mm apart) along with a reasonable clearance. Only cells expressing fluorescent reporters can be imaged in the reflection mode since this device obstructs transmitted light.

Most of the open-ended questions in the field of cell migration require a range of stress from very small values (25 Pa) to moderate values of the order of few kPa (Bao and Suresh, 2003). Over the duration of an experiment (up to a few hours), creep and other time-dependent processes are likely to cause a drop in the compressive load if the plunger is kept stationary (Ahearne et al., 2005). This sets specific requirements for both the sensor and actuator controlling the deformation of the gel.

The system is therefore automated to keep the stress constant within the constraints highlighted above. A mismatch of around 10 Pa in the desired mechanical load is appropriate. Considering a typical gel sample (1% w/v agarose gels, 2 mm thick), 10 Pa change in the mechanical stress corresponds to a plunger displacement of the order of 1 µm. We therefore choose a motorised actuator with a minimum incremental motion of 0.2 *µm*. The load cell must also be able to sense variations of load of the order of 10 Pa. A Futek sensor, LSB200, and a low-cost 16 bit analog to digital convertor (ADC, Adafruit, ADS 1115) connected to a Raspberry Pi computer provide a required range and precision.

When significant changes in the load are applied (typically at the start of an experiment), a slight drift in the coverslip causes a temporary loss of focus. This is largely due to the compliance of the bottom glass coverslip. The timescale of the load application should be long enough to make sure that the cells are tracked during imposition of load, but short enough that the transient response of the cell can be studied. Hence, the stress is ramped up over approximately 10–20 seconds (plunger speed of 40 *µm*/*s*) so that the focus could be manually adjusted during the plunger motion, keeping cells in focus while recording their transient behaviour.

In addition to characterising cell behaviour under compressive load, the same setup can be used to measure the mechanical properties of the gels under which cells migrate. In the next section, we measure the Young’s modulus of different agarose gels and compare the performance to an standard indenter system used for material characterization demonstrating the general application of this system. We then characterise the effect of mechanical load on the migration of cells and quantitatively capture the dynamics of the plasticity in cell migration.

## Results

### Mechanical characterisation of agarose gels

Spherical indentation testing is a standard method to probe the mechanical properties of soft materials (Field and Swain, 1993). Using a spherical indenter as plunger, it is possible to record the force-indentation curve and extract the effective stiffness of the gel in the relevant time-scales used in this experiment. Figure 2A shows the variation of force with distance during an indentation test of an agarose gel. As demonstrated in Figure 2B, the force increases with an indentation distance according to a power law. The Hertz’s contact model provides a suitable fit for such indentation curves and allows us to estimate the young’s modulus (Hertz, 1882).

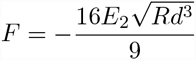

where F is the force applied by an indenting bead, E_2_ is the Young’s modulus of the gel, R is diameter of the bead and *d* is the indentation depth. Figure 2C shows the estimated values of the Young’s modulus for a range of concentrations (0.5%, 0.75%, 1%, 1.5% and 2%). These are effective values as agarose is not a linear elastic material and the apparent stiffness is dependent on the indentation speed due to creep and visco-elastic properties. Nevertheless, we can compare the values obtained with our custom setup with those measured using an industry standard commercial testing machine (Instron 5544) as a validation of our measurement pipeline. The values of Young’s modulus obtained from the cell squasher and the Instron are in good agreement (maximal error of 1.9% for stiffer 2% agarose gel). Thus, it can be concluded that our set-up is capable of applying mechanical stress reliably and reproducibly, and provides a convenient way to measure the Young’s modulus of hydrogels like agarose with sufficient precision.

**Figure 2:**
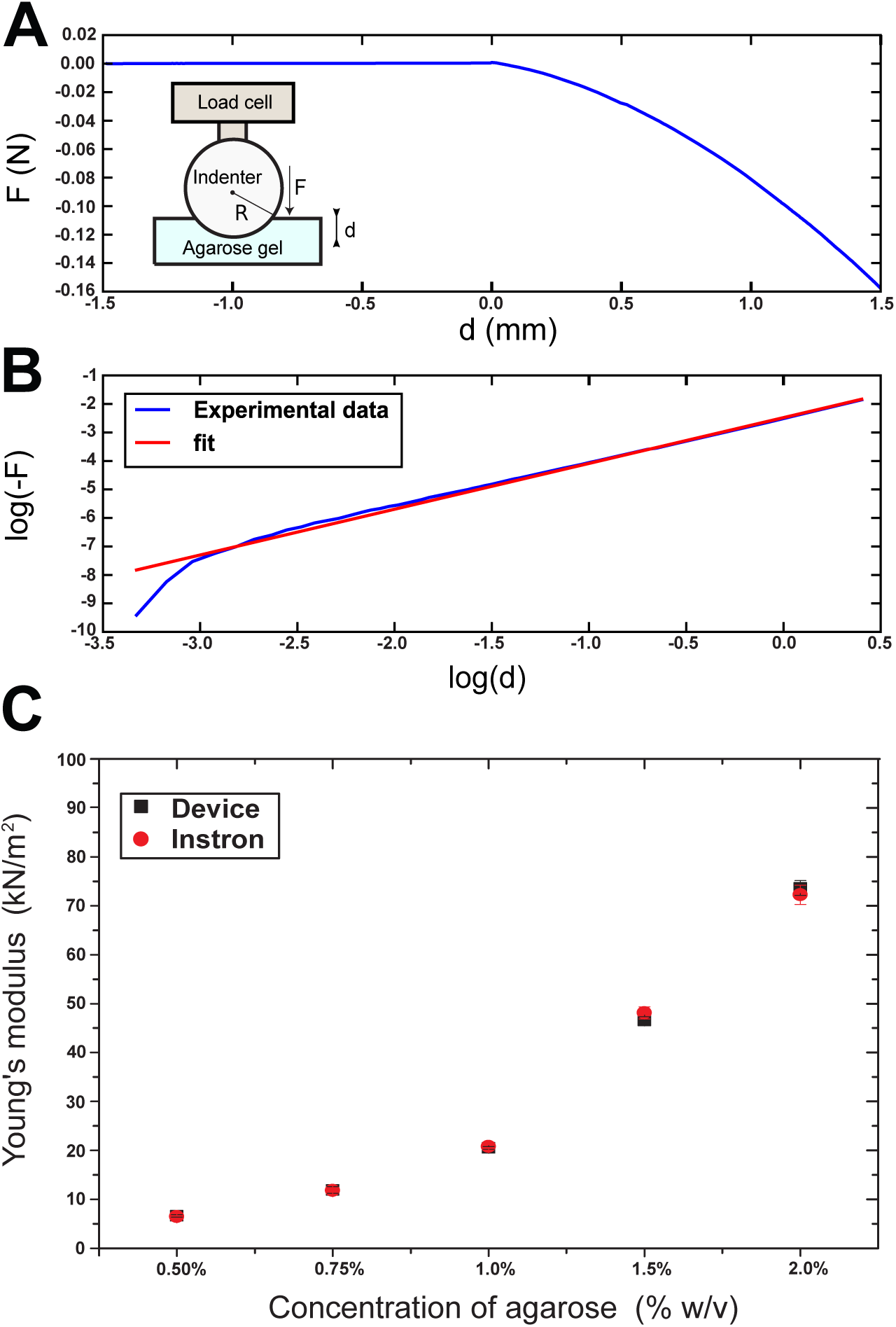
(A) and (B) Force displacement curve for an indentation with a spherical bead of diameter 6 mm on a 1%w/v agarose gel, 5 mm thickness. The indent speed is 0.012 mm/sec. (B) Fit of the force displacement curve with the Hertz contact model: *F* = −*Cd*^3/2^, where *C* =8.38 × 10 ^−2^ *Nmm* ^−3/2^. (C) Young’s modulus of agarose gels of different stiffness measured with the custom device and an Instron 5544 (Canton,MA,USA). For each concentration, two different gels were used, and five different measurements were done on each gel.

Moving back to the flat indenters used to squash the cells under agarose, Figure 3A shows a typical loading curve on a 0.5% agarose gel at 400 Pa load. Initially, the load remains constant as the plunger approaches the agarose gel. The load increases slightly when the plunger comes in contact with the gel due to surface tension between the rectangular plunger and agarose gel surface. It takes approximately 20 seconds for the load to reach the desired value. Thereafter, the device maintains the load at a constant value, within the 10 Pa precision specified above. The device constantly adjusts the position of the plunger, as shown in Figure 3B, to maintain a constant stress by compensating for the creep occurring in the gel. Thus, we demonstrate here that our device can apply a specified load on an agarose gel in a relatively short timescale, and is able to maintain it at a constant value over a long duration.

**Figure 3:**
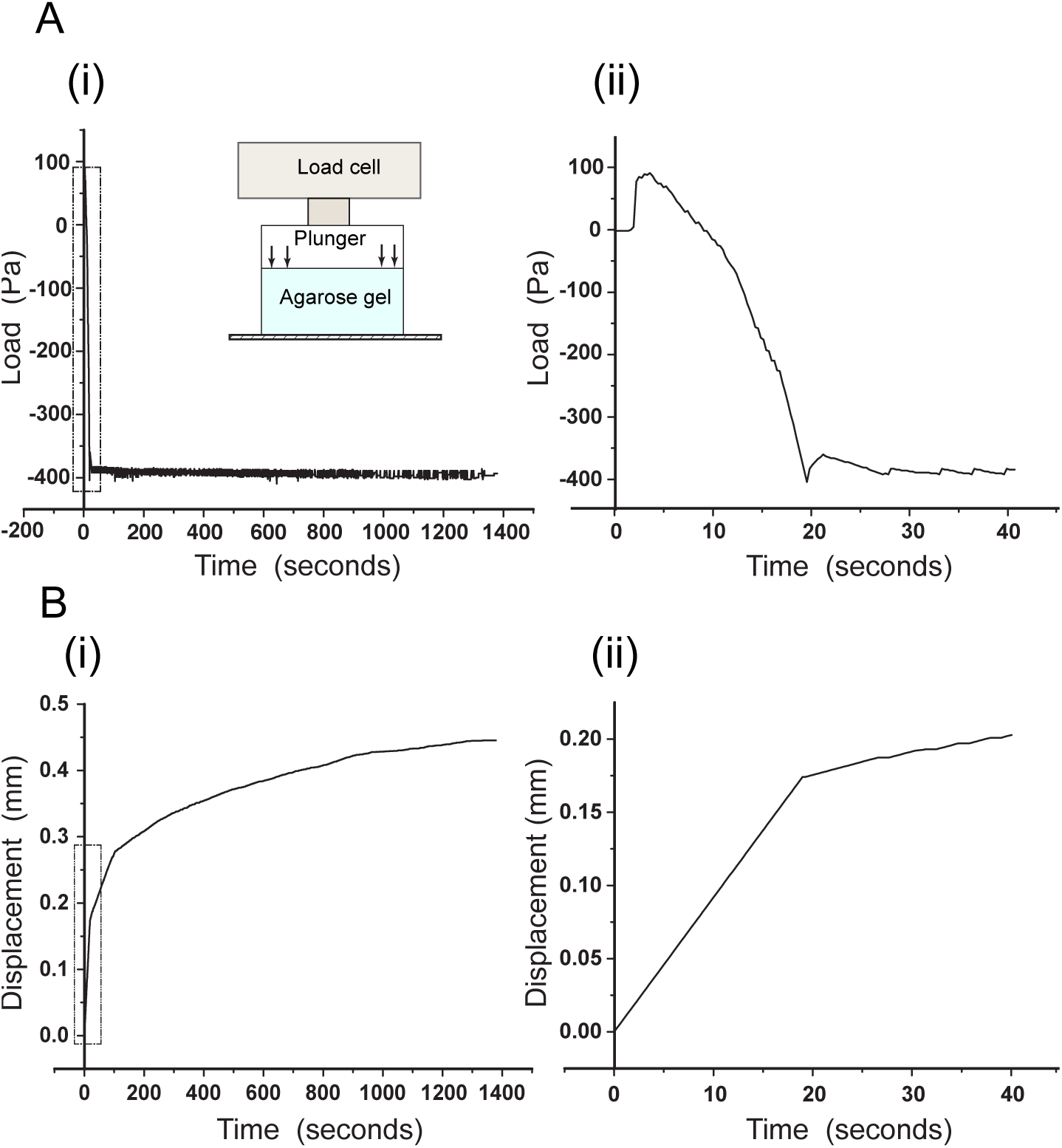
Indentation of agarose gel with flat indenter during cell squashing experiments. (A) Typical loading experiment on a 0.5% agarose gel, 2 mm thickness at 400 Pa load. (i) Variation of load at longer time-scales during a typical experiment (ii) Details of the response during the loading stage. (B) Displacement of the flat indenter for the corresponding loading condition (i) over longer time-scales (ii) during the loading stage.

## Application to biological experiments

### Imposing a compressive load on *Dictyostelium discoideium*

In order to squash cells and probe their response, cells need to be present at an interface between the gel and the glass-bottom dish (see Figure 1A). There are two main approaches to achieve this, as illustrated in Figure 4.

**Figure 4:**
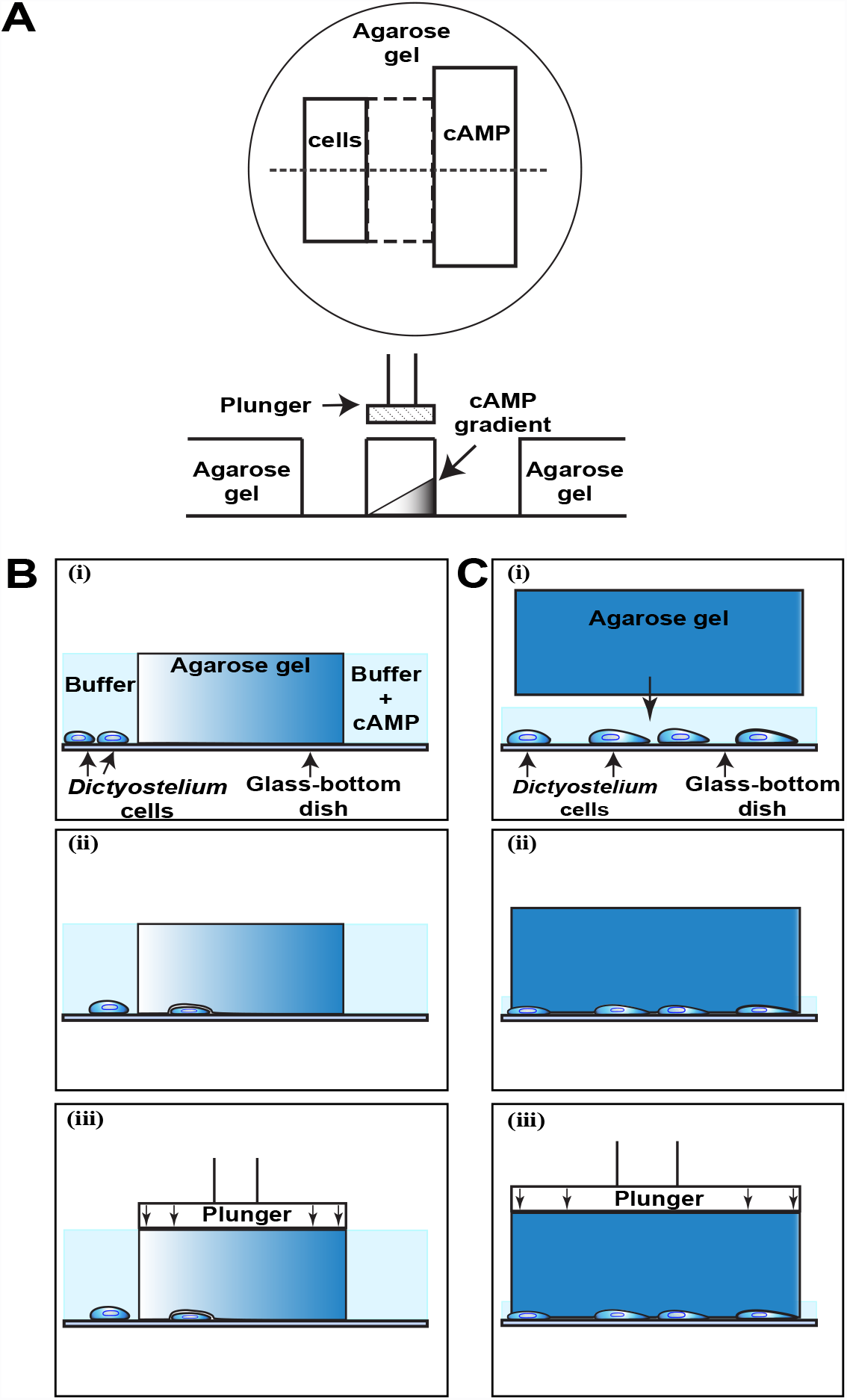
Two mechanisms to place cells under agarose. (A-B) Cells could be attracted underneath an agarose gel by chemoattractant. (C) A layer of agarose could be placed on them.

Chemotactic cells, such as aggregation competent *Dictyostelium* cells, can be persuaded to migrate under the gel using a chemoattractant gradient. A conventional setup, an under agarose assay, involves two independent wells cut in a homogeneous agarose slab of gel, as depicted in Figure 4A. The distance between the wells is 4 mm, and the gel thickness is typically 2 mm. One well is filled with a cell suspension (2 × 10^5^ cells/mL), and the other with a solution of cAMP (5 *µ* M), a strong chemoattractant for *Dictyostelium* cells. After 15-20 minutes, cells start to respond to the chemotactic gradient and migrate under the agarose slab. Figure 4B depicts the protocol used in such a configuration. The experiment starts without any load applied. Once cells are visible on the microscope under the gel, the plunger imposes a set load on the gel’s surface. Alternatively, to probe cells whose migration would not respond to directional cues, cells can be directly squashed by placing a slab of agarose on top of them (see Figure 4C). In both cases, once a load is imposed with the plunger, the gel makes contact with the cells and transmits mechanical forces to them, as demonstrated below.

### Effect of a compressive load on *Dictyostelium discoideum*

To verify that the cells are effectively compressed by the applied load, we measure their height under a range of compressive stresses on them, under 0.5% agarose gel. The height of the cells is quantified by taking z-stacks of *Dictyostelium* cells expressing ABD-GFP which is a reporter for F-actin (Pang et al., 1998). 3-D images were reconstructed and the distance between the top and bottom of the cells was measured. These z-stacks were also corrected for z-axis elongation which occurs as a result of a refractive index mismatch between oil on the microscope lens and aqueous media of the cells (Traynor and Kay, 2007). As seen in Figure 5, the height of the cells decreased from 7 *µm* without any load down to 3 *µm* under 400 Pa. A load of 100 Pa was sufficient to squash the cells by 30%. The height seemed however to plateau for loads beyond 500 Pa. The data therefore demonstrates that the cell squasher operates in a range of forces appropriate to significantly deform the cells and record their morphological evolution.

**Figure 5:**
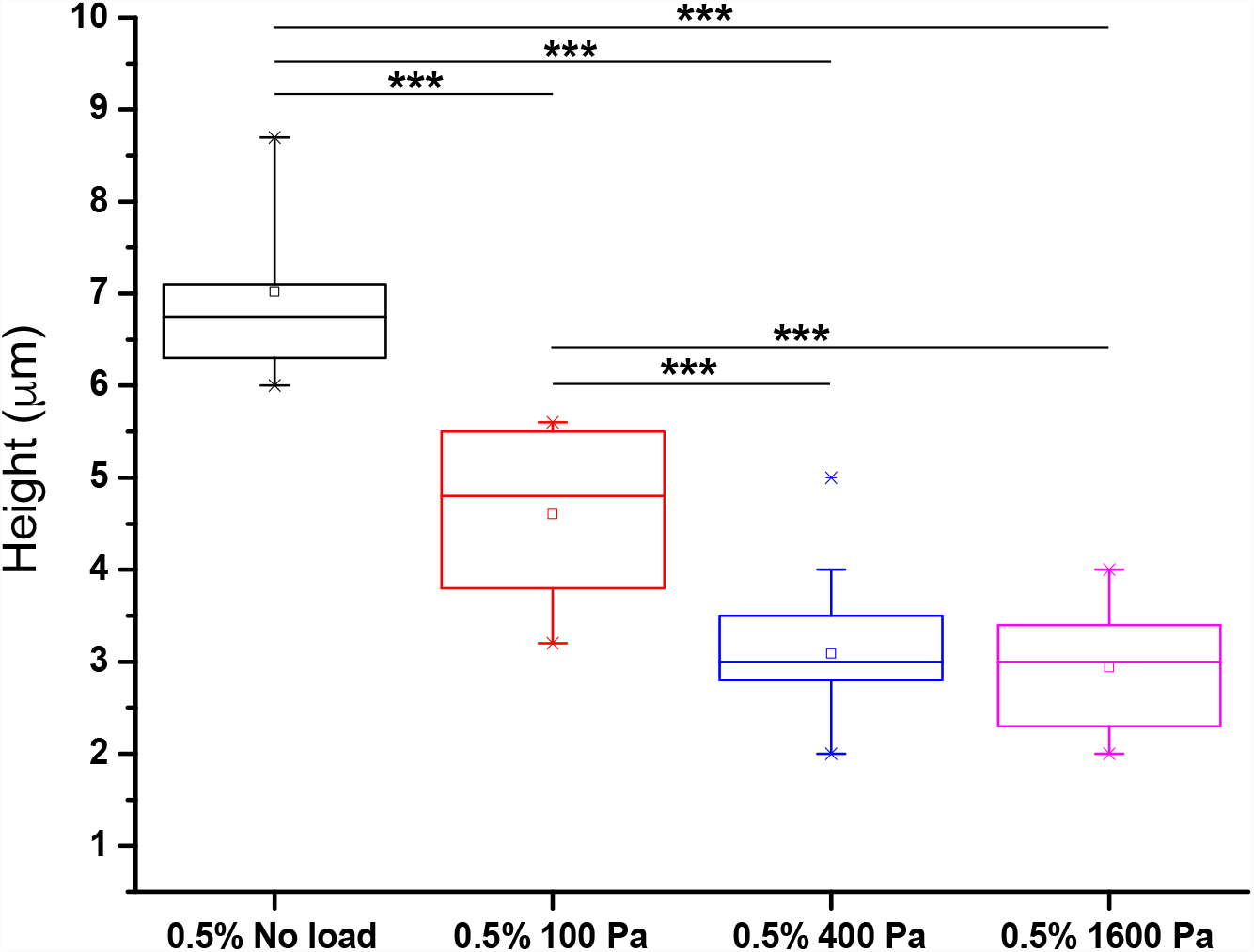
Cell height as a function of the imposed uniaxial load, under a gel of 0.5% agarose (n=20 cells for each case and p<0.001, Tukey’s means comparison test and One Way ANOVA).

### Effect of mechanical stress on the migration of cells

In addition to morphological traits, it is also possible to characterise the behaviour of the cells and in particular their migration under load. As an example, we studied the transition from pseudopods to bleb driven migration. Zatulovskiy et al. showed that *Dictyostelium* cells switch to bleb mode of migration when the stiffness of an agarose layer, underneath which they migrate, is increased from 0.5% to 2%. We first use the chemoattractant based assay introduced in Figure 4B with low concentration (0.5% w/v) agarose. We indeed observe highly polarized cells migrating towards the gradient of cAMP, using pseudopods as their primary mode of migration (Figure 6A). As soon as a mechanical load of 100 Pa is applied, cells tend to form blebs (Figure 6A and Supplementary movie M1). This behaviour is also observed using the alternative protocol highlighted in Figure 4C without cAMP. Initially the cells migrate randomly by forming multiple pseudopods in different directions. The application of a compressive load triggers almost instantly the formation of blebs (see Figure 6B and Supplementary movie M2).

**Figure 6:**
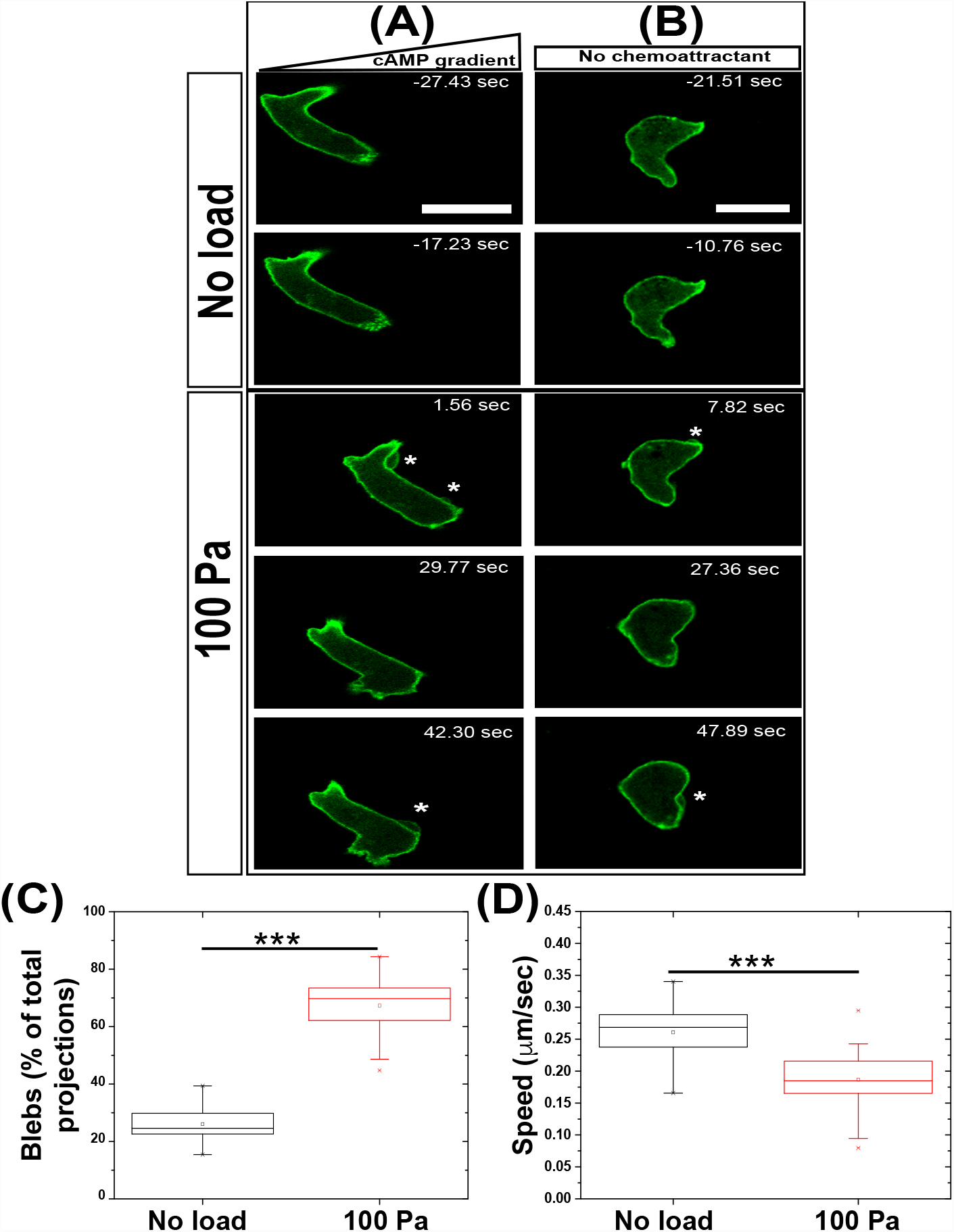
Protrusion formation of cells changes upon dynamic loading. Dynamic behaviour of the cells when migrating (A) along a chemotactic gradient or (B) without a gradient. *Dictyostelium* cells express ABD-GFP. Time indicated on each image is relative to the time at which the plunger makes contact with the 0.5% agarose gel. The imposed load is 100 Pa. Asterisks highlight the location of blebs on each image. (C) and (D) Quantification of Blebs as proportion of total protrusions formed by cells and Speed of migrating cells. (n≥20 cells for each condition and p<0.0001, Tukey’s means comparison test and One Way ANOVA).

We quantified the proportion of blebs amongst total protrusions formed by the cells to establish the effect of compressive load on protrusion formation. Under no load conditions and 0.5% agarose gels, blebs formed 25 ± 6.1 % of total protrusions which increased to 66.8 ± 9.4 % of total protrusions under 100 Pa steady-state load. Along with transition in type of protrusion, the speed of cells also decreased from 0.26 *µm*/sec under no load condition to 0.18 *µm*/sec under 100 Pa load (Figure 6C).

In order to quantitatively analyze the transition observed in Figure 6, a record of both the load sensor signal and timing of bleb events for a particular experiment is presented in Figure 7 (and Supplementary movie M3). The frequency of blebs strongly increases after the load is applied, with a first bleb triggered a few seconds after the compressive load starts to increase. After the load reaches its steady value of 400 Pa, up to 6 blebs are recorded over a minute.

**Figure 7:**
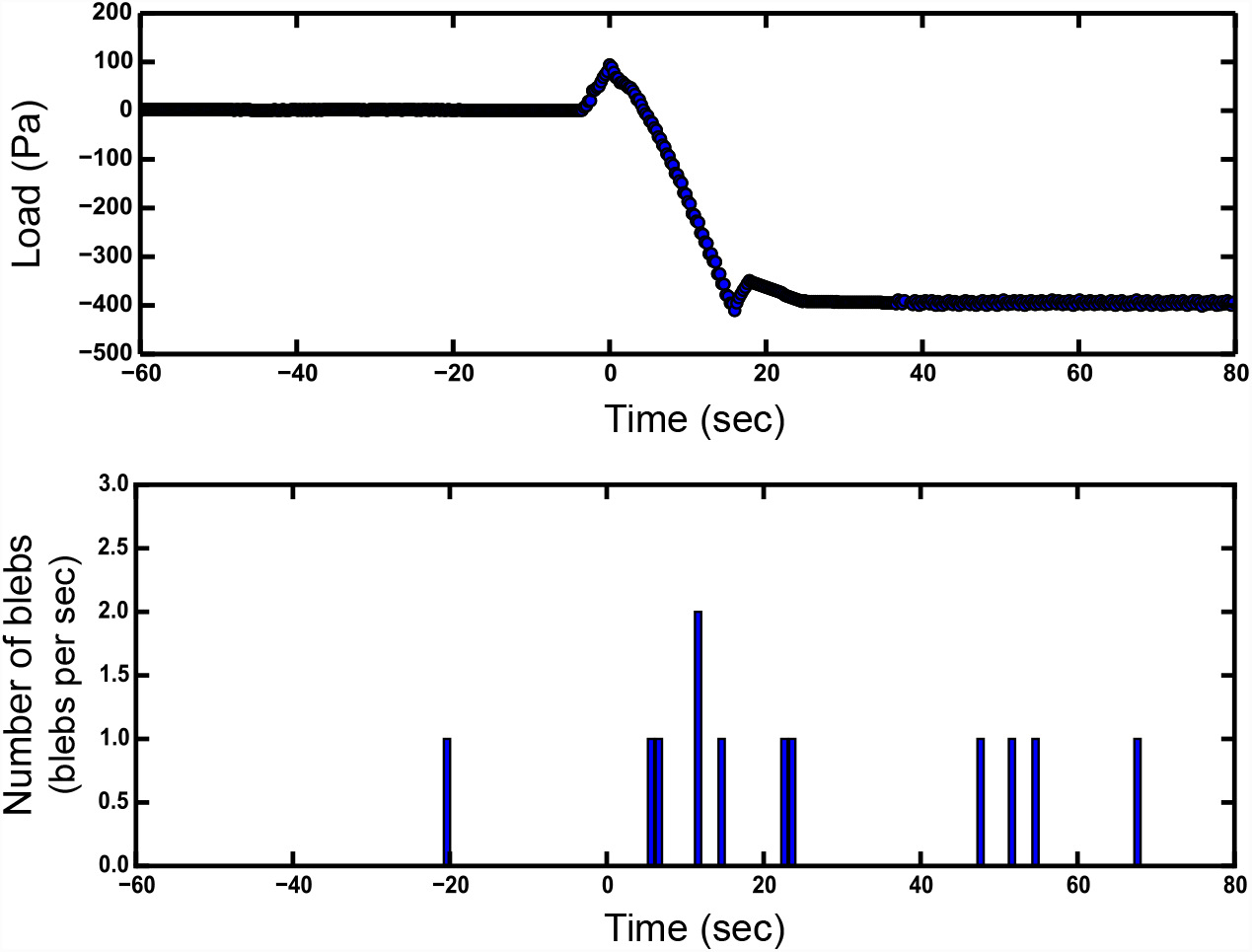
Quantitative analysis of the dynamic response of *Dictyostelium* cells to uniaxial loading. (A) Record of mechanical load as a function of time. t=0 corresponds to the moment when the plunger makes contact with the gel. (B) Occurrence of blebs as a function of time. This is a representative data for a single cell, migrating under 0.5% agarose and corresponding measurement from the cell squasher while applying 400 Pa load.

This establishes that the cell squasher influences the migratory behaviour of cells by the imposition of a mechanical load alone and allows us to quantitatively study the transition.

## Discussion

We have designed an automated device that accurately controls a mechanical compressive load at the interface between a migrating cell and a standard hydrogel such as agarose. Combining mechanical loading with live cell imaging enables us to quantitatively analyze cells’ response to mechanical perturbations. This was demonstrated by monitoring the morphological evolution and migration of *Dictyostelium* cells under compression. As an illustration of the system’s capabilities, we present novel evidence that cells change their migratory behavior within seconds of the imposition of the load, getting flatter and transitioning from pseudopod- to bleb-based migration. Although a chemotactic cue is a convenient way to drive cells under an agarose overlay, similar results are also obtained by simply positioning the gel on top of non-chemotactic cells.

In addition to biological measurement, the same setup also provides a convenient method for characterizing hydrogel’s stiffness and creep behavior over relevant length-scales and time-scales. Young’s modulus values measured on the agarose gels used in the experiments are in excellent agreement with measurements made using an industry standard testing machine that many laboratories might not be able to afford or have access to. One attractive aspect of the setup presented here is therefore to provide a stand-alone rig to characterize both the mechanical properties of the environment and the response of the cells, which a total cost on the order of about three thousand dollars, largely due to the value of the linear stage.

The reason why this system is simple is that the load is applied globally on the gel. This contrasts with more local manipulations, such as interactions with an AFM tip. Here, we do not directly control the exact value of the load experienced by the cells under the agarose. In addition to the applied load, cells are likely to experience stresses due to the elastic deformation of the gel, and probably contribute to the remodeling of gel around them. The setup is however directly designed to control the key parameters of the cell’s mechanical environment, stiffness and load, such that comparison with *in vivo* conditions are possible.

The instrument is also generic enough to be used on other cell types and reduce the dearth of tools which could be utilized to gather a quantitative understanding of mechanosensing response of cells performing their normal physiological functions. *Dictyostelium* cells being amongst the fastest cells, there is no doubt that many other cell types, including mammalian cells, could be studied without additional technical difficulties, besides maybe tuning the range of forces sensed and applied. The set-up would for instance be useful to study the migration of neutrophils in order to gain quantitative understanding about the immune response, or cancer cell invasion under mechanical constraints (Friedl and Alexander, 2011). Coupling such measurements with the activity of mechanosensing pathways, such as YAP/TAZ, is likely to contribute to our understanding of mechanotransduction and its dynamics during individual or collective cell migration.

## Materials and Methods

### Measurement of Young’s modulus of agarose gels

The Young’s modulus of agarose gels was measured by indentation tests. The gels were prepared as 5 mm thick agarose slabs of following concentrations 0.5%, 0.75%, 1%, 1.5% and 2% (%w/v). Indentation tests were performed with the velocity of the bead movement set at 0.012 mm/sec. A round steel bead of diameter of 6 ± 0.01 mm was used as an indenter.

The Young’s moduli of the gels were also measured using a commercially available indentation instrument Instron [Instron 5544 (Canton,MA,USA)]. Spherical indentation tests was performed on these gels to measure their stiffness.

The values were obtained as load v/s extension curves and they were used to calculate the Young’s modulus of the agarose gel using hertz model by considering a non adhesive elastic contact between the agarose gel and the indenting spherical bead (Hertz, 1882).

### Cell culture and Reporters

Experiments were done using the axenic strain Ax2 (Kay laboratory strain; DBS0235521 at http://dictybase.org) of *Dictyostelium discoideum*. Cells were grown at 22°C in HL5 axenic medium in suspension.

*Dictysotelium discoideum* was transformed with a F-actin reporter, ABDGFP to visualize protrusions. This reporter consists of a ABP-120’s F-actin domain (residues 9-248) that are fused to GFP and driven by strong actin-15 promoter (Pang et al., 1998); transformation of Ax2 with this reporter gave strain HM2040. Selection pressure was maintained by growing the cells in the medium containing 10 *µ*g/ml of G-418 antibiotic. This allowed selecting for cells expressing this vector.

### Preparation of cells for under-agarose and squashing experiments

KK2 buffer (16.5 mM KH_2_PO_4_, 3.8 mM K_2_HPO_4_, 2 mM MgSO_4_, and 0.1 mM CaCl_2_) was used as a standard buffer for all the experiments. Aggregation-competent *Dictyostelium* cells were used for these assays. They were prepared by harvesting vegetative cells in exponential phase of growth. 2 × 10^7^cells/ml were resuspended in KK2 and shaken at 180 RPM and 22°C for 1 hour and then pulsed with 90 nM (final concentration after pulsing) cAMP (cyclic-AMP) at every 6 min for further 4.5 hours. Cells stick to the glass walls and form small clumps upon reaching the stage of aggregation competency.

### Microscopy and image analysis

Cells were imaged on a glass-bottom dish (35 mm dish with 10 mm glass bottom) (MatTek corporation) using an inverted laser-scanning confocal microscope (Carl Zeiss, LSM780 or 710) with a 63×/1.4 NA oil immersion objective. Images were collected using Zen 2010 software (Carl Zeiss) and processed using ImageJ (NIH).

Cell height was measured by reconstruction of z-stacks (0.4 *µm* increments) using Imaris (Bitplane). z-axis elongation, partially due to mismatch in refractive indices (Hell et al., 1993), was determined by comparing the z-and x/y-axes of fluorescent beads (9.7 *µm* diameter FluoSpheres; Molecular Probes) and corrected by dividing stack increments by 1.97 (giving true z-axis increments of 0.203 *µm*).

Blebs are fast projections initially devoid of F-actin and were distinguished from the pseudopods using morphological evidence of an actin scar left behind when a bleb is formed. These projections were also distinguished using a kymograph by measuring rate of membrane projections. Number of blebs and pseudopods formed by a cell was counted and the ratio of blebs and total projections was calculated.

The speed of cells was calculated by automated tracking using QUIMP plug-in in Image J (Tyson et al., 2010). The centroid of cells was tracked at every frame of the movie to calculate the total distance covered by the cell. It was then divided by total time to calculate the speed of the cells.

Statistical analysis of the cell height measurement and speed of cells was done using One-way ANOVA and Tukey’s means comparison test in Origin software (Originlabs).

### Cell migration assays

A modified version of the under-agarose assay was used (Laevsky and Knecht, 2001) to attract cells under the agarose as previously mentioned in (Zatulovskiy et al., 2014). Briefly, 0.5% w/v SeaKem GTG agarose (Lonza biochemicals) prepared in KK2 was poured into a preheated glass-bottom dish (35 mm dish with 10 mm glass bottom) to a height of 2 mm. Two parallel rectangular troughs were cut 4 mm apart once the agarose gel has cooled and set. The larger trough was 4 mm wide and 8 mm long while the smaller one was 1 mm wide and 5 mm long. 5 µM cAMP was added and left for about 30 min to allow a gradient of chemoattractant to be set up within the agarose layer, after which 2 × 10^5^cells/mL aggregation-competent cells were placed in smaller trough. The load was applied using the device once the cells started to migrate under an agarose overlay.

The compressive load could also be applied on the cells without subjecting them to an under-agarose assay or chemotactic gradient as shown in Figure 4C. Briefly, a 2 mm thick agarose gel (0.5% w/v) was prepared in a glass-bottom dish. A suspension of 2 × 10^5^aggregation-competent cells/ml was placed in another glass-bottom dish. Once the agarose gel has solidified, it was carefully lifted and placed over the cells. The load was subsequently applied on these cells.

To apply the load using the device, firstly the plunger was carefully brought close to the surface of the gel by manual positioning using the micrometers and then the command to apply the desired load was given. A log file recorded the applied load at all the instances which was used for further data analysis. Quantitative analysis of the number of blebs corresponding to an applied mechanical load was done by measuring the number of blebs in cell migration movies and corresponding data of applied load from the log file of the device.

## Acknowledgments

The authors would like to thank the workshop unit at Department of Engineering and MRC-LMB. The authors are also grateful to N. Barry for help with microscopy and D. Traynor for help and support in assembling the device and incorporating it with the microscope. This work is supported by a Dr. Manmohan Singh Scholarship from St. John’s College to N. Srivastava, Medical Research Council core funding MC_U105115237 to R.R. Kay and BBSRC grant BB/K018175/1 to A.J. Kabla.

